# Reduction in voluntary activation of elbow and wrist muscles in individuals with chronic hemiparetic stroke

**DOI:** 10.1101/689364

**Authors:** Lindsay R. P. Garmirian, Julius P. A. Dewald, Ana Maria Acosta

## Abstract

After a stroke, descending drive is impaired due to the loss of corticospinal and corticobulbar projections which causes a reduction in voluntary activation or an inability of the nervous system to activate muscles to their full capacity, which in turn contributes to weakness of the upper extremity. Voluntary activation has not been quantified at specific joints in the upper extremity, in part because directly assessing changes descending drive is difficult. In this study, voluntary activation of elbow and wrist flexors and extensors was assessed in participants with chronic hemiparetic stroke using twitch interpolation. Twitch interpolation uses electrical stimulation to estimate voluntary activation and relies on the principle that there is an inverse relationship between the amplitude of a twitch evoked by a stimulus and the voluntary force output during stimulation (Taylor, 2009). We measured voluntary activation using twitch interpolation as well as maximum voluntary torque (MVT) of the elbow and wrist flexors and extensors in the paretic and non-paretic limb of ten participants post stroke and the dominant and non-dominant limb of 2 control participants. Results show, MVT interlimb differences were significantly greater for stroke participants compared to control, across muscle groups (p≤0.005). For stroke participants, MVT interlimb differences were significantly greater at the wrist compared to the elbow (P=0.003). Voluntary activation was significantly less in the paretic limb compared to the non-paretic, dominant and non-dominant limbs, across participants and muscle groups (p<0.005 for all four muscle groups). For the stroke participants, the voluntary activation interlimb difference was significantly greater for the wrist muscles compared to the elbow muscles (p<0.005). There was a significant positive correlation (*r* = 0.39, *P* = .022) between each participant’s impairment level, as measured by a hand specific subscore of the Fugl-Meyer Assessment, and the wrist extensor voluntary activation in the paretic limb but the relationship was not significant for the other muscle groups.

## Introduction

Stroke is the leading cause of serious long-term disability in the United States(Go et al., 2013). Approximately 50-70% of stroke survivors experience long-term upper-extremity functional impairments (Faria-Fortini, Michaelsen, Cassiano, & Teixeira-Salmela, 2011). The primary motor deficits that lead to these functional impairments are paresis, loss of independent joint control (Dewald & Beer, 2001; L. C. Miller & Dewald, 2012) and to a lesser extent spasticity (Ellis, Schut, & Dewald, 2017). The loss of corticospinal projections compromises voluntary activation, whereby participants are unable to activate their muscles to their full capacity, which in turn contributes to weakness. The extent of losses in voluntary activation at specific joints in the upper extremity has not been quantified, in part because directly assessing changes in descending drive is difficult. This paper presents results on the quantification of the extent of losses in voluntary activation of the elbow and wrist flexors and extensors in participants with chronic hemiparetic stroke based on the twitch interpolation method.

After a stroke, descending drive is impaired due to the loss of corticospinal neurons. Previous work has shown that this loss of corticospinal neurons after a stroke results in the inability to activate individual motor units (McComas, Sica, Upton, & Aguilera, 1973), decreased firing rates of individual motor units (Rosenfalck & Andreassen, 1980; Tang & Rymer, 1981) and impaired rate coding (L. M. McPherson et al., 2016). While there is evidence for each of these alterations in motor unit activation post hemiparetic stroke, the overall effect on the extent of loss in voluntary activation has not been quantified in the upper extremity.

Twitch interpolation is an experimental method developed to indirectly measure motoneuron drive (Taylor, 2009) and quantify voluntary activation (Klein, Brooks, Richardson, McIlroy, & Bayley, 2010). Voluntary activation is estimated based on measurement of the muscle twitch evoked by an electrical stimulus at rest compared to during volitional muscle contraction (de Haan, Gerrits, & de Ruiter, 2009). This method has been used successfully to show differences in voluntary activation between muscles (Behm, Whittle, Button, & Power, 2002), at different joint angles (Becker & Awiszus, 2001) and in a variety of pathologies including stroke (Klein et al., 2010; Smith et al., 2016) (Horstman et al., 2008; M. Miller, Flansbjer, & Lexell, 2009).

The quantification of voluntary activation using twitch interpolation relies on the principle that there is an inverse relationship between the amplitude of a twitch evoked by a stimulus and the voluntary force output during stimulation (Taylor, 2009). In other words, if an electrical stimulus is applied when a muscle has already been fully activated volitionally, no additional force will be evoked but if a muscle is stimulated at rest, a large amplitude twitch will be evoked. Two different methods are most often employed, the ratio method and the linear prediction method. In the ratio method, voluntary activation is estimated by comparing the amplitude of the twitch elicited by the electrical stimulus applied during rest to the twitch elicited by stimulation during a maximum voluntary torque (MVT). In contrast, in the linear prediction method, voluntary activation is equivalent to the maximum voluntary torque normalized by the theoretical maximum joint torque. In this method, stimulation is applied at rest and during 33%, 66% and 100% of MVT. The theoretical maximum joint torque is obtained by extrapolating the value of volitional torque at which electrical stimulation would no longer elicit a twitch response based on a linear regression of volitional torque versus the measured twitch amplitude. The ratio method was used for this study because most participants post stroke cannot reliably perform wrist flexion and extension contractions at 33% and 66% of MVT.

Voluntary activation has been quantified post stroke during isometric maximal voluntary contractions using twitch interpolation in the lower extremity. More studies have used twitch interpolation in the lower extremity compared to the upper extremity because the neuromuscular anatomy of the lower extremity allows for easier stimulation of the majority of agonist muscles via a peripheral nerve. For example, to measure voluntary activation of the plantar flexors, one can stimulate the tibial nerve to activate all of the primary plantar flexors. Horstrman et al. reported a 57.8% activation of the knee extensors post stroke compared to 93.6% in control limbs (Horstman et al., 2008) while Miller et al. reported 86% activation in the paretic knee extensors versus 97% in the non-paretic knee(M. Miller et al., 2009). Klein et al. reported 48% voluntary activation in the paretic plantar flexors compared to 97% in the non-paretic ankle (Klein et al., 2010). Less twitch interpolation data exists studying voluntary activation following neural injuries in the upper extremity because the neuromuscular anatomy is less amenable to this approach, compared to the lower extremity (Becker & Awiszus, 2001; Fimland et al., 2011; Horstman et al., 2008; Klein et al., 2010; M. Miller et al., 2009; Smith et al., 2015). There is no easily accessible peripheral nerve that can be stimulated to activate all of the primary elbow or wrist flexors or extensors and one must activate individual muscles through stimulation of the motor point instead. While the upper extremity anatomy presents some technical challenges, the technique has been successfully adapted following neural injuries due to spinal cord injury (Peterson et al., 2017) and multiple sclerosis(Sheean, Murray, Rothwell, Miller, & Thompson, 1997). However, to our knowledge, no one has quantified voluntary activation in the upper extremity post stroke. This study is the first to adapt and apply twitch interpolation to both the elbow and wrist flexors and extensors in any population and thoroughly quantify voluntary activation in the upper extremity post stroke.

The goal of this study was to quantify voluntary activation of the paretic and non-paretic upper extremity (elbow and wrist flexors and extensors) in participants post stroke as well as in the dominant and non-dominant upper extremity of able-bodied participants using a twitch interpolation protocol developed by the authors (Garmirian, Acosta, Hill, & Dewald, n.d.). We hypothesized that voluntary activation would be decreased in the paretic limb, compared to the non-paretic limb and both limbs of the able-bodied participants. We also hypothesized that, consistent with previous findings (L. M. McPherson et al., 2016), distal muscles are more affected by stroke and thereby voluntary activation is decreased at the wrist compared to the elbow in the participants post stroke but not in able-bodied participants.

## Methods

### Participants

Voluntary activation and strength of the elbow and wrist flexors and extensors was assessed in the paretic and non-paretic limbs of ten individuals with chronic hemiparetic stroke and in the dominant and non-dominant upper extremity of two control participants (Table 4.1). The Shirley Ryan AbilityLab clinical neuroscience research registry database was used to recruit the ten participants with chronic hemiparetic stroke. All lesions had occurred at least 3.7 years prior to participation (mean ± SD: 11.0 ± 2.5 years, Table 4.1). Participants were excluded if they had a severe concurrent medical problem, an acute or chronic painful or inflammatory condition of either upper extremity or diabetes. The research protocol was approved by the Institutional Review Board of Northwestern University and participants provided written informed consent.

**Table 4.1.** Participant demographics

| Participant | Gender | Group | Affected | Dominant limb | Age (years) | Years Post stroke | Fugl-Meyer Upper-Extremity |
| --- | --- | --- | --- | --- | --- | --- | --- |
| 1 | M | Stroke | R | R | 55.8 | 3.7 | 24 |
| 2 | F | Stroke | R | R | 63.1 | 8.0 | 30 |
| 3 | M | Stroke | L | R | 45.6 | 5.4 | 33 |
| 4 | F | Stroke | R | R | 60.4 | 12.8 | 25 |
| 5 | M | Stroke | R | R | 65.4 | 8.5 | 18 |
| 6 | F | Stroke | L | R | 61.6 | 28.1 | 16 |
| 7 | M | Stroke | R | R | 69.4 | 19.0 | 12 |
| 8 | M | Stroke | R | R | 63.5 | 14.7 | 48 |
| 9 | M | Stroke | L | R | 62.2 | 5.0 | 47 |
| 10 | M | Stroke | R | R | 44.6 | 5.0 | 38 |
| **Mean (SD)** |  |  |  |  | **59.2 (2.6)** | **11.0 (2.5)** | **9.1 (4.0)** |
| 11 | F | Control | NA | R | 64.1 |  |  |
| 12 | F | Control | NA | R | 53.4 |  |  |

### Isometric torque measurements

Participants were seated in a Biodex chair (Biodex, Shirley, NY) with their trunk secured via shoulder and lap belts and the tested arm rigidly attached to an isometric setup to measure elbow and wrist joint torques. A 6 degree-of-freedom load cell (JR3, Woodland, CA) was used to measure elbow flexion and elbow extension torque (Dewald & Beer, 2001) while a single axis torque sensor (Futek, Irvine, CA) mounted at the wrist was used to measure wrist torque. The participant was positioned with 15° shoulder flexion, 30° shoulder abduction, 90° elbow flexion and in neutral with respect to pronation/supination and wrist and finger posture.

### Twitch Interpolation Setup

A standard twitch interpolation protocol (Behm, St-Pierre, & Perez, 1996; Dowling, Konert, Ljucovic, & Andrews, 1994; Peterson et al., 2017) was used with the following specifications and modifications to optimize measurement of the paretic upper extremity. Using the locations listed in Table 4.2 as starting points, a motor point pen (Compex, Guildford, UK) was used to determine the specific location of each motor point for each muscle by finding the point where the lowest amplitude of stimulation triggered the largest visible muscle contraction (Cheong, Hong, & Chung, 2013; B. K. Park, Shin, Ko, Park, & Baek, 2007; Perotto, 2011). Two self-adhesive electrodes (Dura-Stick, Hanover, Germany) were placed on each muscle group, one over the specific motor point (cathode) and the other distal to the motor point (anode) as specified in Table 4.2.

**Table 4.2.** Electrode Location and Size

| **Muscle Group** | **Approximate Motor Point Location (Cathode)** | **Distal Electrode Location (Anode)** | **Electrode Size** |
| --- | --- | --- | --- |
| Elbow flexors | 4 Finger breadths distal to axilla, 1 cm anterior to medial bicipital groove (short head of biceps brachii) | Over distal biceps tendon | 5cm x 6cm |
| Elbow extensors | 5 finger breadths from posterior axilla along a line from the posterior axilla to the olecranon (long head of triceps) | Over distal triceps tendon | 5cm x 6cm |
| Wrist extensors | 2 finger breadths distal to lateral epicondyle of humerus, along a line from the lateral epicondyle to the base of 2<sup>nd</sup> metacarpal (extensor carpi radialis longus and brevis) | Bony prominences of posterior wrist | 4cm x 6cm |
| Wrist flexors | 3 finger breadths distal to medial epicondyle, along a line from the medial epicondyle of the humerus to the base of the 2<sup>nd</sup> metacarpal (flexor carpi radialis) | 2 finger breadths proximal to anterior wrist crease | 4cm x 6cm |

### Maximum Voluntary Torque Measurements

For each of the four muscle groups, participants were asked to generate at least six maximum voluntary torques (MVTs) in each degree of freedom (elbow flexion, elbow extension, wrist flexion and wrist extension); additional trials were collected if needed to ensure “true” maximum voluntary torque, by restricting the six greatest torque values to be within 10% of each other and to guarantee that the last trial did not result in the greatest value. Participants were provided with real time visual feedback of torque performance as well as verbal encouragement to produce maximum contractions.

### Twitch Interpolation Measurements

The electrical stimulus consisted of a single monophasic pulse with duration of 100μs to maximize motor recruitment while minimizing activation of pain fibers. The maximal stimulation amplitude was set at the 1mA increment where the measured joint torque plateaued or began to decrease. This is in contrast with standard twitch interpolation protocols in the lower limb that employ 110% of the maximal stimulation amplitude (Klein et al., 2010). Due to the reduced muscle and limb volume in the upper-extremity compared to the lower extremity, using supramaximal amplitudes will result in current spread to antagonist muscles, decreasing net joint torque production.

Participants were asked to relax followed by a maximum voluntary contraction held for 4 seconds (Fig. 4.1). A stimulus of maximal amplitude was applied after 250 ms of sustained torque level within 10% of the target effort. A second stimulus was applied 6 s after the first stimulus, with the limb at rest (Fig. 4.1). This was repeated for 6 trials. Between trials, participants were asked to activate the antagonist muscle group, to eliminate residual torque induced by persistent muscle activation from the previous trial (J. G. McPherson, Ellis, Heckman, & Dewald, 2008). This is an important consideration since individuals post stroke often have hypertonicity and difficulty relaxing their muscles, especially the flexor muscles, thus the use of reciprocal inhibition is needed to minimize any potential residual torques from the previous trial.

**Figure 4.1.**
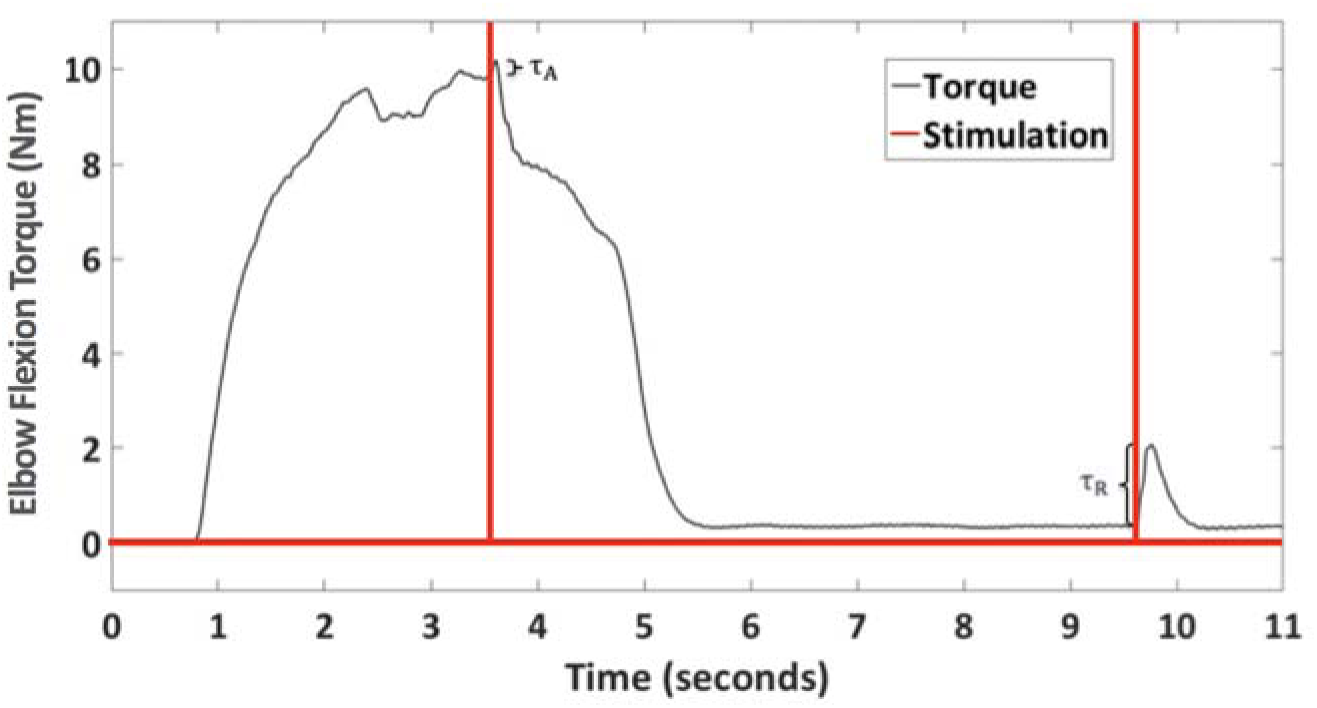
Typical trial data from an elbow extension twitch interpolation trial for the paretic limb of one of the chronic stroke participants. Electrical stimulation, shown in red, has been superimposed on the elbow extension torque data, shown in black.

### Data Analysis

Elbow joint torque was calculated offline from the measured forces and moments at the forearm through a Jacobian-based transformation. The wrist torque was obtained directly from the torque sensor based on its calibration equation determined previously. All joint torques were filtered using a two-sided moving average filter with a 100 ms window. For the maximum voluntary torque (MVT) trials, MVT was calculated as the greatest average torque maintained over 0.5 seconds. For stroke participants, MVT interlimb difference (%) was calculated as the difference in torque generated between the non-paretic and paretic limbs, normalized by the non-paretic limb. For control participants this was calculated as the difference between the dominant and non-dominant limbs, normalized by the dominant limb.

For each of the four muscle groups, voluntary activation was estimated for each trial using the formula defined in (1):

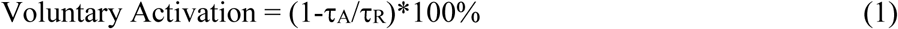

Where τ_A_, the twitch torque, represents the additional torque produced by electrical stimulation during MVT and τ_R_, the resting twitch torque, represents the torque produced by electrical stimulation with the limb at rest (Fig. 4.1) (Allen, Gandevia, & McKENZIE, 1995). The pre-stimulus torque value was defined as the average torque produced over the 250 ms window prior to the stimulus. The stimulus torque was defined as the maximal torque value elicited within 150 ms after the stimulus. Therefore, τ_A_ and τ_R_ represent the net torque generated by electrical stimulation during a MVT and rest, respectively. Note that 100% voluntary activation indicates that the participant is able to fully activate the tested muscles, whereas 0% indicates complete loss of volitional control of the tested muscles. For stroke participants, voluntary activation interlimb difference (%) was calculated as voluntary activation of the non-paretic minus the paretic. For control participants this was calculated as the dominant minus the non-dominant.

### Statistics

To test if MVT interlimb difference was different in the participants post stroke versus control participants, we used a repeated measures mixed model with group (control, stroke) as a fixed factor. To test if MVT interlimb difference was greater at the wrist compared to the elbow within the stroke participants, we used a repeated measures mixed model with joint (elbow, wrist) as the fixed factor. To test if percent voluntary activation was different in the paretic, non-paretic, non-dominant or dominant limb for all four muscle groups, we used a repeated measures mixed model with limb (paretic, non-paretic, non-dominant or dominant limb) and muscle group (elbow flexion, elbow extension, wrist flexion, wrist extension) as fixed factors. To test if voluntary activation interlimb difference was different at the elbow versus the wrist within the stroke participants, we used a repeated measures mixed model with joint (elbow, wrist) as the factor. A Bonferonni correction was used to account for multiple comparisons where appropriate. Pearson correlation coefficients were used to determine if there was a significant correlation between voluntary activation and impairment level as measured by a subscore of the upper-extremity Fugl-Meyer Assessment that only includes the wrist and hand components (sections VI and VII). A p-value of ≤0.05 was considered significant.

## Results

### Typical trial data

Fig. 4.1 shows example data from an elbow extension twitch interpolation trial for the paretic limb of one of the stroke participants. Electrical stimulation, shown in grey, has been superimposed on the elbow extension torque data, shown in black. The percent activation for this trial was calculated as 48%.

### Maximum Voluntary Torque

The average MVT for the paretic limb was 41.6, 33.2, 9.12 and 6.48 Nm for the elbow flexors, elbow extensors, wrist flexors and wrist extensors respectively and 69.7, 50.8, 25.8 and 15.8 Nm for the non-paretic limb (table 4.3). The average MVT for the non-dominant limb of control participants was 49.0, 33.4, 22.9 and 10.6 Nm for the elbow flexors, elbow extensors, wrist flexors and wrist extensors respectively and 50.0, 32.4, 23.9 and 15.8 Nm for the dominant limb (Table 4.3). MVT interlimb difference was significantly greater for stroke participants compared to control, across muscle groups (p≤0.005), as illustrated in Figure 4.2. For the stroke participants, the MVT interlimb difference was significantly greater at the wrist compared to the elbow (P=0.003), as seen in Fig. 4.2.

**Table 4.3.**
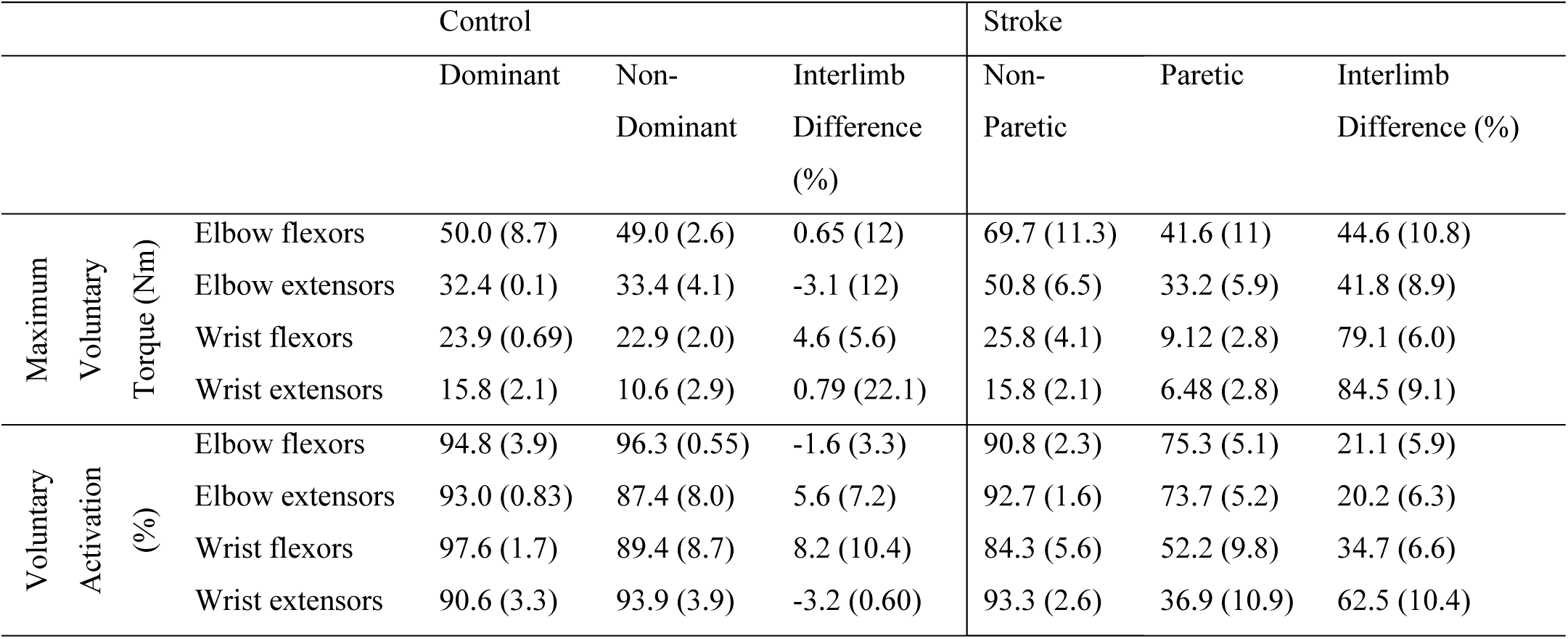
Mean (standard error) maximum voluntary torque (MVT) and voluntary activation for the dominant and non-dominant limbs of the control participants and non-paretic and paretic limbs of the stroke participants as well as the average interlimb difference for MVT.

|  |  | Control |  |  | Stroke |  |  |
| --- | --- | --- | --- | --- | --- | --- | --- |
|  |  | Dominant | Non-Dominant | Interlimb Difference (%) | Non-Paretic | Paretic | Interlimb Difference (%) |
| Maximum Voluntary Torque (Nm) | Elbow flexors | 50.0 (8.7) | 49.0 (2.6) | 0.65 (12) | 69.7 (11.3) | 41.6 (11) | 44.6 (10.8) |
|  | Elbow extensors | 32.4 (0.1) | 33.4 (4.1) | -3.1 (12) | 50.8 (6.5) | 33.2 (5.9) | 41.8 (8.9) |
|  | Wrist flexors | 23.9 (0.69) | 22.9 (2.0) | 4.6 (5.6) | 25.8 (4.1) | 9.12 (2.8) | 79.1 (6.0) |
|  | Wrist extensors | 15.8 (2.1) | 10.6 (2.9) | 0.79 (22.1) | 15.8 (2.1) | 6.48 (2.8) | 84.5 (9.1) |
| Voluntary Activation (%) | Elbow flexors | 94.8 (3.9) | 96.3 (0.55) | -1.6 (3.3) | 90.8 (2.3) | 75.3 (5.1) | 21.1 (5.9) |
|  | Elbow extensors | 93.0 (0.83) | 87.4 (8.0) | 5.6 (7.2) | 92.7 (1.6) | 73.7 (5.2) | 20.2 (6.3) |
|  | Wrist flexors | 97.6 (1.7) | 89.4 (8.7) | 8.2 (10.4) | 84.3 (5.6) | 52.2 (9.8) | 34.7 (6.6) |
|  | Wrist extensors | 90.6 (3.3) | 93.9 (3.9) | -3.2 (0.60) | 93.3 (2.6) | 36.9 (10.9) | 62.5 (10.4) |

**Figure 4.2.**
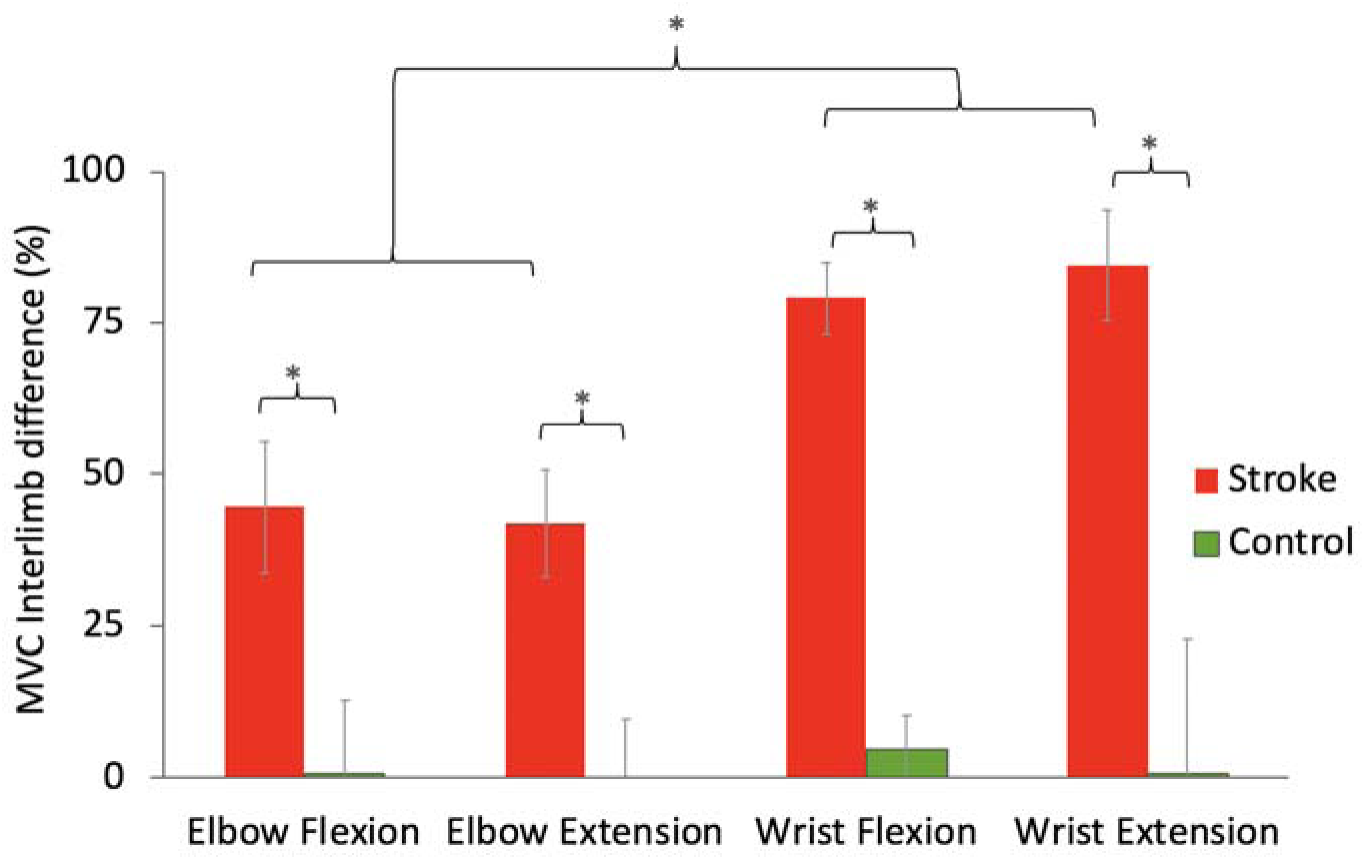
Maximum voluntary torque interlimb difference for stroke and control participants for the elbow and wrist flexors and extensors. For stroke participants this was calculated as: (non-paretic – paretic)/non-paretic and for the control participants this was calculated as (dominant - non-dominant)/dominant. Results are the average ± one standard error for ten stroke participants and two control participants.

### Voluntary activation

Voluntary activation was significantly smaller in the paretic limb compared to the non-paretic, dominant and non-dominant limbs, across participants and muscle groups (p<0.005 for all four muscle groups) (Fig. 4.3). The average voluntary activation for the non-paretic limb was 94.8, 93.0, 97.6, 90.6% Nm for the elbow flexors, elbow extensors, wrist flexors and wrist extensors respectively and 71.7, 72.5, 48.7 and 26.4% for the respective paretic limb muscle groups (Table 4.3). There was no significant difference in voluntary activation between the dominant limb, non-dominant and non-paretic limb, as seen in Fig. 4.3 (p=1.0). The average activation for the dominant limb of control participants was 94.8, 93.0, 97.6 and 90.6% for the elbow flexors, elbow extensors, wrist flexors and wrist extensors respectively and 96.3, 87.4, 89.4 and 93.9% for the respective muscle groups in the non-dominant limb (Table 4.3). For the stroke participants, the activation interlimb difference was significantly greater for the wrist muscles compared to the elbow muscles (p<0.005), as seen in Table 4.3.

**Figure 4.3.**
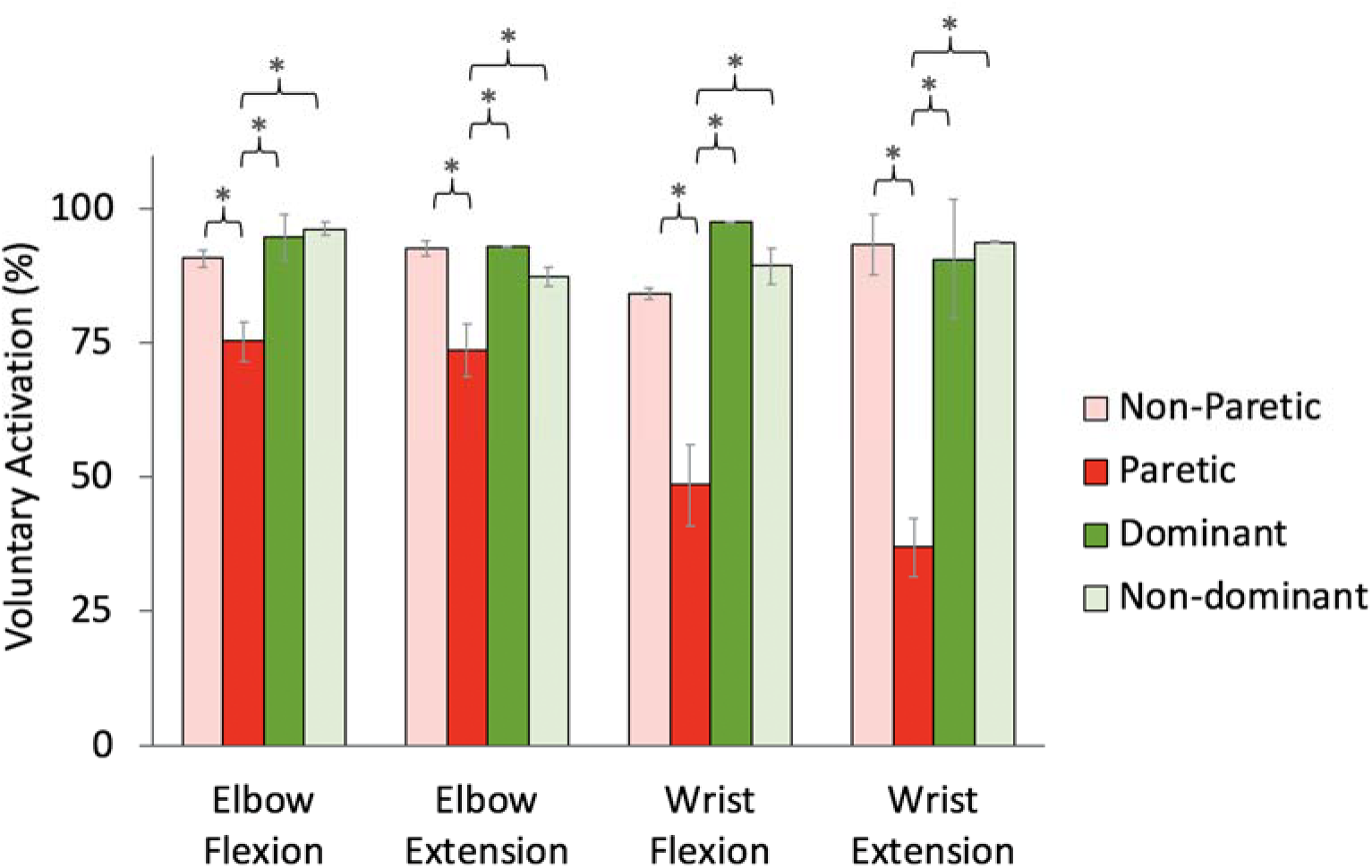
Percent voluntary activation for the non-paretic and paretic limb of stroke participants (red) and the dominant and non-dominant upper extremity of control participants (green) for the elbow flexors and extensors and wrist flexors and extensors. Results are the average ± one standard error for the ten stroke participants and two control participants.

### Impairment level versus voluntary activation

There was a significant positive correlation (*r* = 0.39, *P* = .022) between each participant’s impairment level, as measured by the hand subscore of the Fugl-Meyer Assessment, and the wrist extensor percent activation (Fig. 4.4). Interestingly, this relationship was not significant when correlating Fugl-Meyer (or the hand specific subscore), with activation of the elbow flexors (*r* = 0.52, *P* = .119), elbow extensors (*r* = 0. 46, *P* = .185) or wrist flexors (*r* = 0.40, *P* = .519).

**Figure 4.4.**
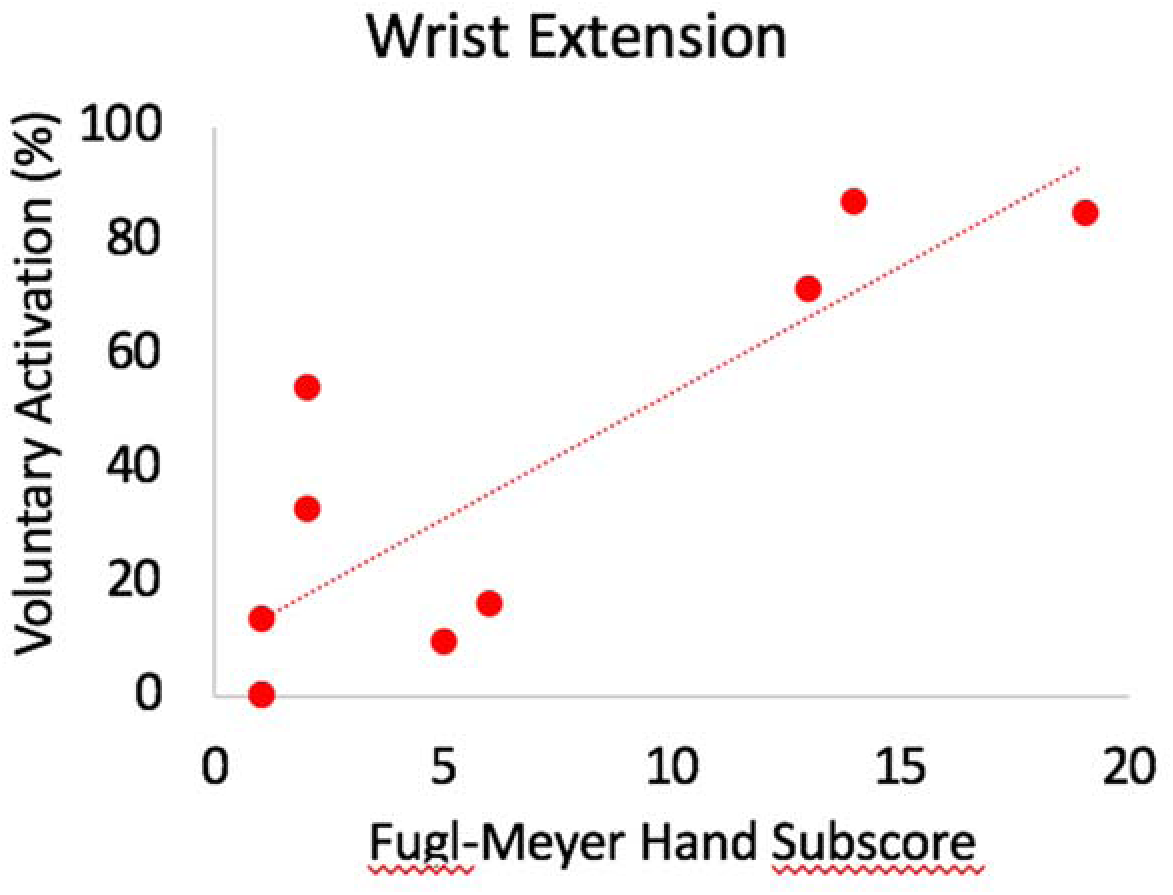
Wrist extensor percent voluntary activation as a function of paretic upper extremity Fugl-Meyer Assessment hand and wrist score (sections VI and VII). Each data point represents a single participant post stroke. Pearson correlation analysis indicates a significant positive correlation between percent activation and impairment level (*r* = 0.84, *p* = .02).

## Discussion

This study presents estimates of voluntary activation of the elbow and wrist flexors and extensors in participants post stroke as well as control participants. This is the first time that the extent of losses in voluntary activation has been reported in the upper-extremity post stroke. We found that voluntary activation was significantly decreased in all muscle groups of the paretic limb, compared to the non-paretic and control limbs. Additionally, we found that voluntary activation was more impaired distally at the wrist, compared to the elbow, in participants post stroke.

The results of this study agree with previous studies that show decreased voluntary activation in the lower-extremity post stroke. Past studies have reported voluntary activation in the paretic lower extremity of ambulatory participants post stroke of 57.8% in the knee extensors (Horstman et al., 2008), 48% in the plantar flexors (Klein et al., 2010) and 86% in the knee extensors (M. Miller et al., 2009). Our results indicate that voluntary activation of proximal muscles in the upper extremity (elbow flexors: 71%, elbow extensors 72%) may be less impaired compared to the lower extremity (although one study found 86% activation in the lower extremity), while voluntary activation of distal upper extremity muscles may be more impaired compared to the lower extremity (wrist flexors: 48%, wrist extensors: 26%). However, these comparisons between studies must be made cautiously, due to important differences in the protocols and setups used to determine voluntary activation.

Several factors contribute to decreased voluntary activation after stroke, including inability to activate individual motor units(McComas et al., 1973), decreased firing rates of individual motor units(Rosenfalck & Andreassen, 1980; Tang & Rymer, 1981)and impaired rate coding (L. M. McPherson et al., 2016). All of these factors will decrease the efficacy of activation and the ability to produce joint torque. Additionally, greater deficits in rate coding at distal muscles (L. M. McPherson et al., 2016) may explain why voluntary activation is more impaired at the wrist compared to the elbow.

Interestingly, there was a significant positive correlation between impairment level and wrist extensor percent activation but not with other muscle groups. This means that participants post stroke with greater impairments had less voluntary activation at their wrist, compared to participants with less impairment. This is interesting, given the evidence that shows deficits in wrist and finger extension are early predictors of recovery post stroke (Nijland, van Wegen, Wel, & Kwakkel, 2010). We hypothesize that this correlation exists because more impaired participants may have greater damage to their corticospinal tract compared to less impaired participants which could cause a decrease in voluntary activation (Stinear et al.; Puig et al.; Schaechter et al.). While the other muscle groups exhibited a similar pattern when correlated with impairment level, it was not statistically significant because of lack of power with the study sample size and limited voluntary activation range within the measurement sample.

Twitch interpolation is a valid method to indirectly measure the degree of voluntary activation after stroke; however, interpretation of the results needs to be constrained by the actual measurements. Twitch interpolation estimates voluntary activation based on torque measurements during voluntary contraction compared to the torques elicited during electrical stimulation. Twitch interpolation does not directly measure descending drive, motoneuron firing rate or the source of motoneuron drive(Taylor, 2009). The method relies on the assumption that the amplitude of the voluntary torque and the amplitude of the evoked twitch are linearly related (inversely)(Taylor, 2009). However, linearity is a simplification of this relationship that ignores tendon compliance, the length-tension relationship, antidromic collisions and stimulation of antagonists. As described previously(Garmirian et al., n.d.), steps were taken to optimize the isometric measurements including minimizing spillover to antagonist muscles while optimizing stimulation of agonists via careful selection of stimulation properties, electrode size and placement.

This is the first study, to our knowledge, to estimate voluntary activation in the upper-extremity post stroke using twitch interpolation. Our study confirms that there is a significant decrease in voluntary activation in participants with chronic stroke and that these deficits are greater at the wrist compared to the elbow. These deficits in voluntary activation have a direct impact on joint strength and can contribute to difficulty completing ADLs. The methods described in this paper can be used to investigate therapeutic interventions that allow individuals with stroke to increase their muscle voluntary activation and address post-stroke weakness.

